# A chemically inducible muscle ablation system for zebrafish

**DOI:** 10.1101/2022.04.20.488934

**Authors:** Eric Paulissen, Benjamin L. Martin

## Abstract

An effective method for tissue specific ablation in zebrafish is the *nitroreductase/metronidazole* system. Expressing bacterial *nitroreductase (ntr)* in the presence of nitroimidazole compounds causes apoptotic cell death, which can be useful for understanding many biological processes. However, this requires tissue specific expression of the *ntr* enzyme, and many tissues have yet to be targeted with transgenic lines that express *ntr*. We generated a transgenic zebrafish line expressing *ntr* in differentiated skeletal muscle. Treatment of embryos with metronidazole caused muscle specific cell ablation. We demonstrate this line can be used to monitor muscle regeneration in whole embryos and in transplanted transgenic cells.

Summary statement

## INTRODUCTION

The study of muscle regeneration is an active field of study with broad applications for therapeutics (Yu et al., 2021). Zebrafish in particular have a broad capacity for regeneration of both the heart and skeletal muscle, and are a useful organism for studying both topics (Beffagna, 2019; Pipalia et al., 2016; Ratnayake et al., 2021; Seger et al., 2011; Wang et al., 2011). In order to faithfully study muscle regeneration, tissue-specific methods of injury are critical to induce a precise biological response to that injury. One method of doing this is by performing targeted ablation of muscle tissue.

Ablation of specific tissues is an important tool for understanding developmental biology. Ablation has been used in understanding topics ranging from regeneration to tissue interactions (Curado et al., 2007; Patterson and Parichy, 2013; Wang et al., 2011). In the zebrafish, common methodologies for tissue ablation include laser ablation, resection, and chemical ablation (Curado et al., 2008; Otten and Abdelilah-Seyfried, 2013; Wang et al., 2011). A common system, the *metronidazole/nitroreductase* system (*MTZ/ntr*), utilizes a bacterial enzyme that reduces nitroimidazole compounds into potent DNA cross linkers (Anlezark et al., 1992; Bridgewater et al., 1995; Curado et al., 2007; Edwards, 1993). The advantage of this system compared to laser or resection methods is the high tissue specificity and temporal control of ablation. Laser and resection methods often damage adjacent tissues, confounding the results and making interpretation difficult. The MTZ/*ntr* system avoids this by only targeting cells that express the *ntr* transgene. However, one disadvantage of this is the availability of transgenic lines that express *ntr* enzymes in the tissue of interest. Here, we describe a transgenic zebrafish line, *tg(actc1b:epNTR-mcherry-caax)*, hereafter referred to as *actc1b:epNTR*, that drives a codon optimized NTR enzyme in the trunk skeletal and heart cardiac muscle.

## RESULTS

### Treatment of transgenic *actc1b:epNTR* zebrafish embryos with MTZ results in ablation of muscle cells

To determine if *actc1b:epNTR* indeed ablated the muscle cells of the zebrafish. Fluorescent microscopy imaging of *actc1b:epNTR* treated with DMSO vehicle showed normal striated muscle fibers in the somites (Figure 1A). In contrast, *actc1b:epNTR* embryos treated with 10 mM MTZ for 30 hours, and washed of MTZ for 24 hours, showed loss of striated muscle fibers and proper somite morphology (Figure 1B). To determine if the loss of muscle fibers is due to cell death, we stained embryos immediately after ablation with acridine orange, a marker of apoptotic cell death in the zebrafish embryo (Lecoeur, 2002; Tucker and Lardelli, 2007). Embryos treated with DMSO vehicle had no acridine orange staining in the somites, showing no detectable cell death in this region of the embryo (Figure 1C). Ablated embryos showed robust staining in the zebrafish somites, indicating broad cell death of the muscle fibers (Figure 1D).

**Figure 1:**
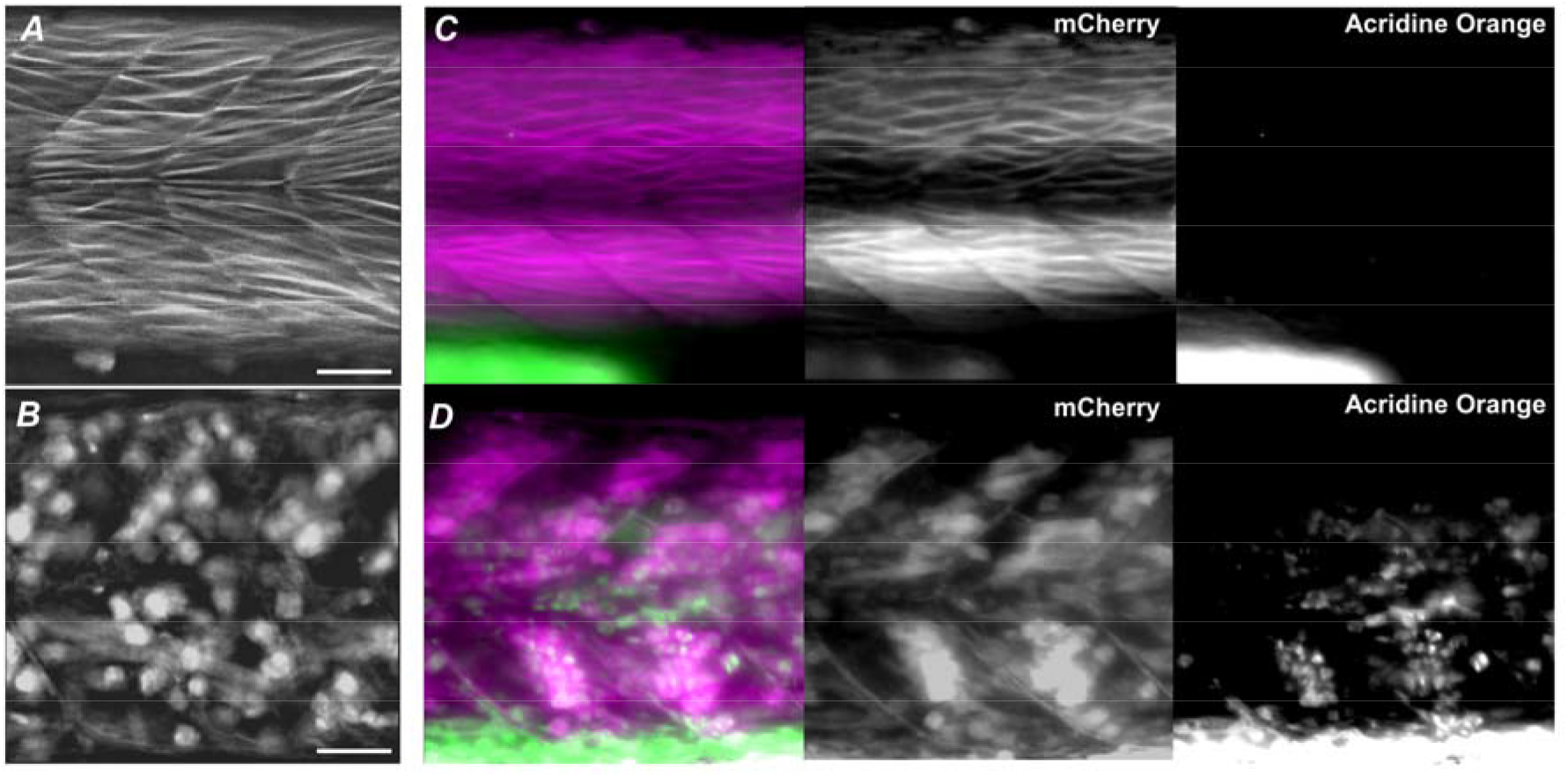
The *actc1b:epNTR* transgene can ablate muscle tissue. (**A, B**) 3 dpf *actc1b:epNTR* embryos treated with DMSO vehicle (A) or 10 mM MTZ (B) at 1 dpf. (**C, D**) 3 dpf *actc1b:epNTR* embryos treated with DMSO vehicle (C) or 10 mM MTZ (D) and stained with 10 μM acridine orange. mCherry and Acridine Orange channels are split to observe lack of acridine orange staining in the DMSO treated embryo. Scale Bars=50 μM

### *actc1b:epNTR* embryos regenerate their musculature after ablation

We investigated whether *actc1b:epNTR* embryos were able to regenerate muscle fibers post-ablation. We imaged embryos over a time course of 96 hours to analyze muscle ablation and regeneration. Embryos at 24 hpf were ablated for 24 hours, at which MTZ was washed out with serial washes in embryo media. We define the time immediately after washing out MTZ as time 0. At time 0, somitic tissue showed numerous rounded cells as well as brightly fluorescent cellular debris (Figure 2A, red and gold arrows, respectively). At 24 hours, the rounded cells have largely disappeared and cellular debris and some striated cells have appeared (gold and white arrows, respectively) (Figure 2B). At 72 hours, many striated cells have begun to form and some cellular debris remains (Figure 2C). At 96 hours, numerous striated cells have regenerated and little cellular debris remains (Figure 2D).

**Figure 2:**
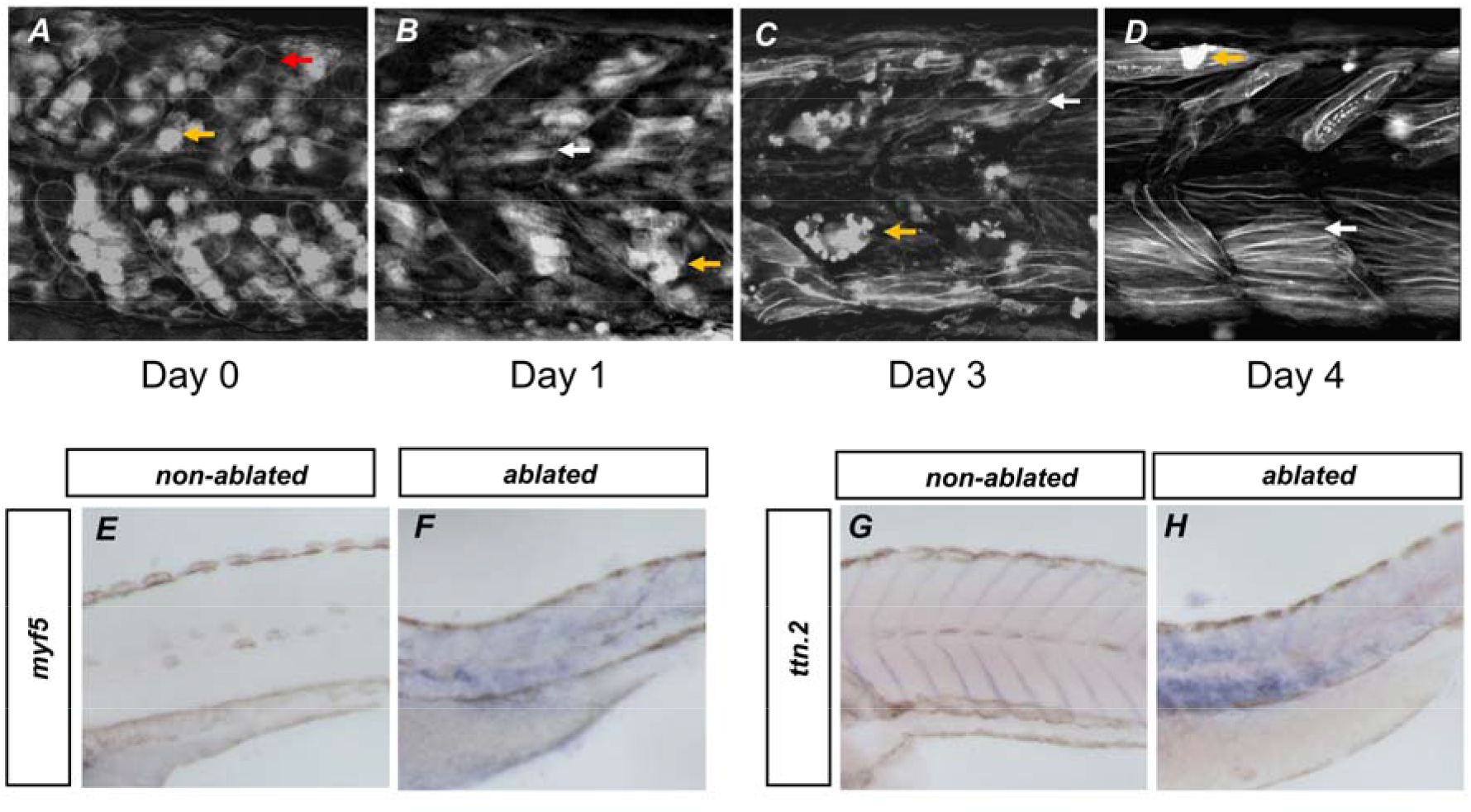
Ablated muscle tissue regenerates over time. (**A-D**): *actc1b:epNTR* embryos ablated with 10 mM MTZ at 1 dpf for 30 hours followed by washout. (**A**) *actc1b:epNTR* embryos imaged for mCherry immediately after washout. (**B**) *actc1b:epNTR* embryos imaged 1 day after washout. (**C**) *actc1b:epNTR* embryos imaged 3 days after washout. (**D**) *actc1b:epNTR* embryos imaged 4 days after washout. (**E-F**): In-situ hybridization for *myf5*. (**E**) Non-ablated embryos. (**F**) Ablated embryos. (**G-H**): In-situ hybridization for *ttn.2*. (**G**) Non-ablated embryos. (**H**) Ablated embryos.

To determine whether the emergence of striated cells indicated a reentrance of regenerative cells into myogenesis, we performed in-situ hybridization against the myogenic marker *myf5* and the muscle marker *ttn.2* (Lin et al., 2006; Yan et al., 2002). This marks both myoblasts as well as differentiated muscle, respectively. In-situ hybridization of *myf5* shows upregulation of *myf5* 72 hours post ablation, whereas in control embryos *myf5* is not expressed (Figure 2E and 2F). In addition, *ttn.2* shows upregulation throughout the somite region after ablation, while non-ablated embryos shows normal expression patterns (Figure 2G and 2H). This indicates that after muscle cells are ablated, cells reenter the myogenic program and generate differentiated muscle fibers.

### Cell death in *actc1b:epNTR* embryos attracts macrophages

A potential benefit of the ablation system is the ability to monitor how different cell types interact with ablated cells. Recent studies have shown how injured muscle tissue interacts with the early immune system, specifically macrophages. Macrophages have been shown to interact with and facilitate myogenesis post injury, as well as clear cellular debris (Dort et al., 2019; Ratnayake et al., 2021). We investigated whether ablated cells in the epNTR model attract macrophages to the ablation site and whether specific macrophage interactions occurred. We generated chimeric embryos by transplanting cells from a *actc1b:epNTR* or wild-type embryo into a *tg(mpeg1:GFP)* host that labels macrophages. We then treated the host embryos with MTZ at 24 hpf for 24 hours. Host embryos containing wild type cells show no strong presence of the macrophages at the location of the transplanted cells, and the macrophages were mainly transient (Figure 3A, Supplemental Fig. 1, Figure 3 Supplemental Video 1). Host embryos that contain *actc1b:epNTR* cells show broad recruitment of macrophages to the site of cell ablation (Figure 3B). In addition, macrophages appear to interact with and intake debris from adjacent cells (Figure 3 Supplemental Video 2). This indicates that dying cells from MTZ/*ntr* system attract the immune system.

**Figure 3:**
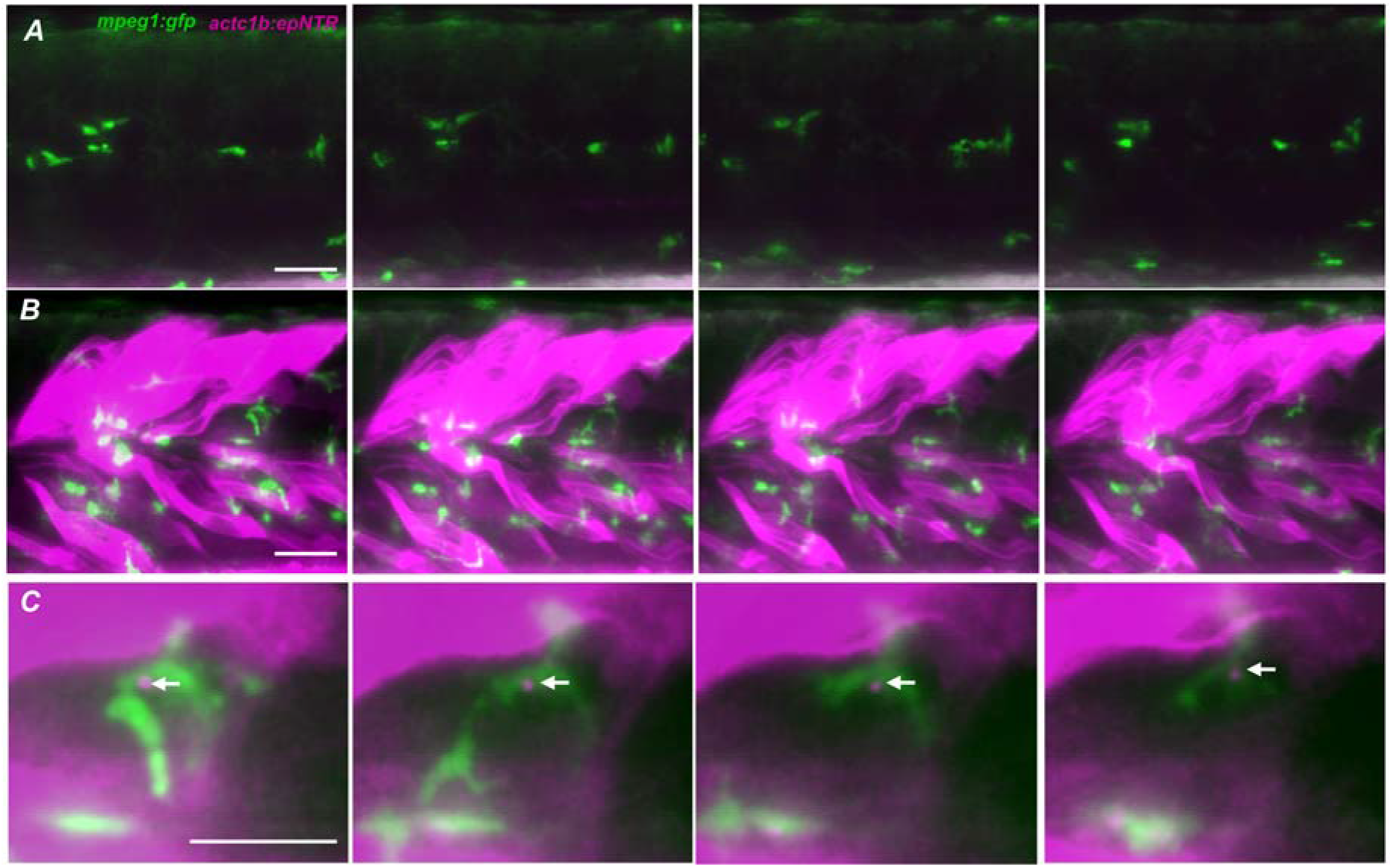
Macrophages interacts with and uptakes apoptosing cells. (**A**) A *mpeg1:GFP* host embryo containing transplanted wild-type cells that lack the *actc1b:epNTR* transgene shows macrophages (green) that do not linger within muscle tissue. (**B**) A *mpeg1:GFP* host embryo with transplanted *actc1b:epNTR* muscle cells (magenta) treated with MTZ and imaged 1 day after washout. Embryos show increased residence of macrophage cells in ablated area. (**C**) A *mpeg1:GFP* macrophage in an *actc1b:epNTR* embryo. mCherry puncta indicates uptake of ablated muscle debris from injured cells (white arrows). Scale Bars=50 μM

We also observed that macrophages take up portions of dying cells. Time-lapse imaging of macrophages show interaction with cells mid-ablation (Figure 3C). Macrophages contain RFP puncta that indicate they have engulfed the epNTR-mCherry-caax fusion protein (Figure 3C, White Arrows). This contrasts with control embryos, in which macrophages show no preference for any tissue (Figure 3A). Macrophages also rarely have RFP puncta in non-ablated embryos relative to ablated.

## DISCUSSION

Here we demonstrate a system which can induce muscle damage within the zebrafish embryo. By utilizing muscle specific *nitroreductase* expression, we can induce cell death specifically within these cells of the zebrafish embryo. An advantage to this system is that ablation is dependent on the drug MTZ, which can be washed out to allow for the embryo to recover. This provides a straightforward methodology for analyzing muscle regeneration within the embryo. Transplantation can also be used to introduce small numbers of muscle fibers expressing *nitroreductase* in an otherwise wild-type host embryo, which allows for ablation of small numbers of muscle fibers in specific targeted regions in the zebrafish musculature.

This particular transgenic line reveals the robustness of the MTZ/*ntr* system. The EpNTR is fused with mCherry-caax in this transgene, and is visibly localized to the cell surface (Figure 1A). However, this does not affect the ability of the fusion protein to ablate muscle tissue. Previously generated transgenic zebrafish utilized cytoplasmic localization of the NTR protein during ablation (Curado et al., 2007; Patterson and Parichy, 2013; Pipalia et al., 2016; Tabor et al., 2014). Evidence that a membrane bound NTR fusion protein can still robustly ablate tissues implies that NTR is not required in a soluble cytoplasmic form to induce cell death.

This system also displays interesting dynamics of cell death and regeneration within the muscle. The advantage of the membrane bound fluorophore in this system is that it allows the visualization of cell structure of both the ablated cells as they are apoptosing and regenerating muscle tissue. Regenerating muscle tissue appears to lose the anterior-posterior polarity that muscle tissue initially adopts in the wild-type zebrafish somite, with some cells unable to span the width of an individual somite. Transplanted NTR positive muscle cells in a wild type environment appear to lose integrity as they begin to undergo apoptosis, adopting a wave-like structure as opposed to straight striations. These observations imply structural deficiencies of dying muscle tissues can be studied using this system. We believe this system is a valuable resource to share with the community.

## MATERIALS AND METHODS

### Generation of the zebrafish transgenic line

The *actc1b:mcherry-caax* plasmid was described previously, and was used as a destination vector for *epNTR* (Berger et al., 2014). The *actc1b:mcherry-caax* plasmid was modified to add an additional EcoRI site and remove the *mcherry* start codon with the two primers GCTCCACCGTGGTGAGCAAG and CCGAATTCCAAACTTGCCTTGGTCTGTGC, both of which had a 5’ phosphorylation modification (Eurofins). Following the PCR amplification using Phusion High-Fidelity DNA Polymerase (NEB), the plasmid was ligated using T4 DNA ligase (NEB).

The *epNTR* gene was obtained through Addgene (Plasmid #62213). The *epNTR* ORF fragment was amplified through Taq PCR using the primers ATGGATATTATTAGTGTGGCC CTG and CCCACCTCTGTCAGTGTGATGTTC. *epNTR* was cloned into a *pCRII* plasmid using the TOPO TA CLONING kit (ThermoFisher). The *PCRII-epNTR* and *actc1b:mcherry-caax* plasmids were cut using EcoRI and the respective DNA fragments were purified by gel extraction (NEB). The isolated *ntr* gene and cut *actc1b:mcherry-caax* were then ligated together using T4 ligase (NEB) to generate the *actc1b:epNTR-mcherry-caax* plasmid.

### Microinjections and strain generation

One-cell stage TLB zebrafish embryos were injected with 25 picograms of *tol2* mRNA along 25 picograms of *actc1b:epNTR-mcherry-caax* plasmid. The *tol2* mRNA was generated as mentioned previously (Goto et al., 2017). F0 embryos were grown to adults and outcrossed to wild-type fish to screen for germline insertion of the transgene. Successful insertions were grown to adulthood.

### Cell transplantation

Donor *tg(actc1b:epNTR-mcherry-caax)* embryos were injected with1 nL of 2% cascade-blue dextran (Invitrogen) as a tracer. Roughly 25-50 cells from sphere stage donor embryos were transplanted into the vegetal pole of host embryos using a CellTram Vario (Eppendorf). Embryos were grown to 24 hpf and were screened by fluorescence for donor cell contribution to the somitic tissue.

### Microscopy and imaging

Fluorescent images of were obtained using a Leica DMI6000B inverted microscope at 20× magnification. Images were processed in ImageJ for transplants or using the Leica Thunder application for whole embryo fluorescence.

### Zebrafish tissue ablation

Metronidazole (Sigma) powder was added directly to embryo growth media (150 mM NaCl, 5 mM KCl, 13 mM CaCl ·2H2O, 1.5 mM KH2PO4,0.5 mM NaHPO4, 10 mM MgSO4 · 7H2O, 1% DMSO) to a working concentration of 10 mM to generate ablation media. The *tg(actc1b:ntr-mcherry-caax)* embryos were grown to 24 hpf and incubated in ablation media at 33°C for 30 hours, unless otherwise notated. Ablation media was then washed out 3 times with embryo media and the embryos were further incubated at 28°C.

### Zebrafish Lines

All wild type embryos used in this study were from hybrid adults generated from an inbred strain of locally acquired pet store fish (which we call Brian) crossed to the TL line (TLB). The *tg(actc1b:epNTR-mcherry-caax)^sbu^* transgenic strain was maintained on the TLB background. The *tg(mpeg1:EGFP)^gl22tg^* line was maintained as homozygotes (Ellett et al., 2011).

## Acknowledgements

We thank Stephanie Flanagan for excellent fish care.

## Competing interests

The authors declare no competing or financial interests.

## Funding

This work was supported by a National Institutes of Health training grant [T32 GM008468] to E.P., and National Science Foundation [IOS1452928] and NIGMS [R01GM124282] grants to B.L.M.

---

**Supplemental Figure 1:**
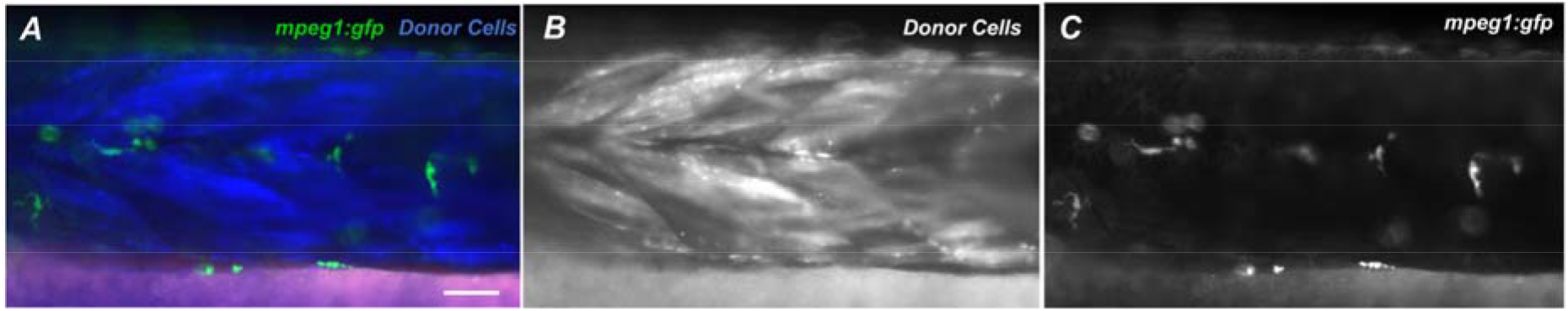
Chimeric wild-type embryos do not show prolonged localization of macrophages. Cascade-blue dextran was injected into wild-type donor embryos and transplanted into *tg(mpeg1:EGFP)* embryos. (**A**) An image showing donor cells in a *tg(mpeg1:EGFP)* embryo. Green indicates macrophages. Blue indicates transplanted cells in the somites. Magenta (absent) indicates lack of expression of a *tg(actc1b:epNTR-mcherry-caax)* transgene. (**B**) Single blue channel from merged image in A. (**C**) Single green channel from merged image in A.

**Supplemental Video 1: Live imaging of macrophages in control embryo**

Live images showing macrophages (green) migrating through the trunk of a 48 hpf zebrafish embryo. Note the lack of directionality or localization of macrophage migration.

**Supplemental Video 2: Live imaging of macrophages during localized ablation**

Live images showing macrophages (green) migrating through the trunk of a 48 hpf zebrafish embryo. Magenta indicates transplanted cells in the trunk of the embryo. Note increased activity of macrophages around the ablating embryos

## Notes

### Competing Interest Statement

The authors have declared no competing interest.

